# Megaevolutionary dynamics in reptiles and the role of adaptive radiations in evolutionary innovation

**DOI:** 10.1101/2020.04.22.055061

**Authors:** Tiago R. Simões, Oksana Vernygora, Michael W. Caldwell, Stephanie E. Pierce

## Abstract

Adaptive radiations are long believed to be responsible for the origin of phenotypic diversity and new body plans among higher clades in the fossil record. However, few studies have assessed rates of phenotypic evolution and disparity across broad scales of time to understand the evolutionary dynamics behind the origin of major clades, or how they relate to rates of molecular evolution. Here, we provide a total evidence approach to this problem using the largest available data set on diapsid reptiles. We find a strong decoupling between phenotypic and molecular rates of evolution, with many periods of accelerated phenotypic evolution or expansion of phenotypic disparity at the origin of major reptile clades and body plans that do not correspond to periods of adaptive radiation. We find heterogeneous rates of evolution during the acquisition of similarly adapted functional types, and that the origin of snakes is marked by exceptionally high evolutionary rates.

---

The classical theory of adaptive radiation predicts that such events are characterized by high rates of phenotypic evolution, in combination with an expansion in phenotypic disparity and taxonomic diversity, as new species rapidly transforming to occupy available adaptive zones during times of ecological opportunity^1,2^. Across geological timescales, lineages undergoing exceptionally fast evolutionary rates would give rise to many new lineages, with some of these fast-evolving lineages potentially going extinct. Once niches are occupied, phenotypic disparity stabilizes and species diversification and evolutionary rates decrease and stabilize at lower levels^1^. It has long been assumed that the aftermath of mass extinctions would provide the ideal ecological opportunities for adaptive radiations^1,3^, such as the diversification of placental mammals after the Cretaceous-Palaeogene mass extinction (KPME), or the appearance of several reptile lineages in the fossil record following the Permian-Triassic mass extinction (PTME)^2,3^. Therefore, adaptive radiations have long been hypothesized to be responsible for the origin of most of biological diversity (in both taxonomic and phenotypic terms), especially regarding the origin of higher clades (e.g. families or orders) and new body plans, or what Simpson had originally referred to as “mega-evolutionary” processes^4^.

Although the concept of adaptive radiations is fundamental to our understanding of evolutionary theory, only recently have quantitative tools been developed to rigorously test its predictions at broad taxonomic and deep time scales in the evolutionary paleobiology. For instance, using both relaxed clocks and phylogenetic comparative methods various studies have found high rates of evolution at the origin of major clades, including the early evolution of birds, arthropods, and crown placental mammals^5–7^. Fast evolutionary rates during putative periods of adaptive radiations following mass extinctions have also been recovered, such as the radiation of birds ^8^ and placental mammals^6^ after the KPME, and archosaurs after the PTME^9^. Overall, these results support the central pillars of adaptive radiation theory concerning the origin and early radiation of major clades.

Nevertheless, there have been important recent challenges to this classical model of adaptive radiation^1,2^. It has been suggested the pattern of fast rates of evolution (“early burst” model) does not seem universal as it was not recovered during the origin and initial radiation of some major groups, such as teleost fishes or echinoids^10,11^. Importantly, very few studies include information from the fossil record and thus cannot examine such megaevolutionary events at sufficiently large scales of time in order to be able to fully comprehend the expected long-term dynamics, events before and after mass extinctions, or the origin of major clades (for notable exceptions, see^11–13^). When observed at broad scales of time, what may appear to be early bursts at the origin of major clades represent episodic events of rapid evolution (episodic radiations) throughout evolutionary history^11^. Additionally, there may exist long macroevolutionary gaps between the origin of clades and new body plans (associated with evolutionary rates and phenotypic disparity) and the actual period of taxonomic diversification (constructive radiations)^14,15^. Therefore, the question of whether adaptive radiations are truly responsible for the origin of most of biological and phenotypic diversity in the history of life and the origin of new body plans and phenotypic novelty remains open.

Here, we explore megaevolutionary dynamics on phenotypic and molecular evolution during two fundamental periods of reptile evolution: i) the origin and early diversification of the major lineages of diapsid reptiles (lizards, snakes, tuataras, turtles, archosaurs, marine reptiles, among others) during the Permian and Triassic periods, as well as ii) the origin and evolution of lepidosaurs (lizards, snakes and tuataras) from the Jurassic to the present. The first provides answers concerning the origin of some of the most fundamental body plans in the history of reptile evolution, as well as the impact of the largest mass extinction event in the history of complex life (the Permian-Triassic Mass Extinction). The second reveals fundamental clues towards the evolution of one of the most successful vertebrate lineages on Earth today, comprising over 10,000 different species^16^. Our major questions for those two chronological and taxonomic categories include: What are the major deep time evolutionary patterns concerning evolutionary rates and phenotypic disparity? Do most periods of expansion of evolutionary rates and/or morphological disparity occur at the origin of major clades and new body plans? correspond to periods of adaptive radiation as predicted by the Simpsonian model? What periods can we identify as conforming to the classical model of adaptive radiation?. Our findings indicate that several periods of reptile evolution undergoing fast evolutionary rates do not conform to the expectations of a classical model of adaptive radiation; and that phenotypic novelties that have converged on similar functions may evolve at distinct rates of evolution.

## Results

We expanded upon our recently published phylogenetic data set of early evolving diapsid reptiles and lepidosaurs (fossils and living)^17^, by adding new data on extant lizards and snakes to inform both phenotypic and molecular components of the tree. To estimate evolutionary rates in a well calibrated evolutionary tree, we integrated both phenotypic and molecular data using total-evidence dating (TED). This is a powerful approach in which tree topology, divergence times and phenotypic and molecular evolutionary rates, are jointly estimated. To account for potential variations in estimates of divergence times and evolutionary rates due to different software implementations, we conducted analyses using the the software Mr. Bayes^18^ and the BEAST2 evolutionary package19. However, the BEAST packages lack diversity sampling strategies, which is known to potentially overestimate divergence times with TED^20,21^. Our results with BEAST2 had relatively older divergence times compared to Mr. Bayes, especially among older nodes, which we attribute to this factor. For all our trees, see Supplementary Information (Supplementary Figs. S1-14 and Supplementary Data).

In our analyses we found evidence for deep root attraction (DRA)^22^ that, when corrected (following^22^), increased the precision for divergence times in Mr. Bayes (Supplementary Figs. S4,5,14), and were also in much greater agreement with the fossil record—e.g. the divergence time for the diapsid-captorhinid split at the earliest Pennsylvanian (322 MYA), thus being close to the age of oldest known diapsid reptiles from the Late Pennsylvanian^23^. In contrast, even in analyses in which we tried to correct for DRA in BEAST2, the median age for the diapsid-captorhinid split was placed at the latest Devonian close to the Devonian-Carboniferous boundary (ca. 40 million years older), a time at which the first known tetrapods were diversifying onto land^24^, and thus, considerably more inconsistent with the fossil record (Supplementary Figs. S6,7). Overestimated divergence times are likely to affect estimates of evolutionary rates by extending chronological branch lengths. Therefore, our results and conclusions are primarily driven from the posterior tree estimates obtained from Mr. Bayes (results from BEAST2 are in our Supplementary Information).

Our initial non-clock Bayesian inference results with lepidosaurs indicate strong topological similarity between our molecular tree and a recent phylogenomic studies of lepidosaurs^25^, especially concerning the paraphyly of amphisbaenians in both instances. Amphisbaenian paraphyly was also obtained by analyzing phenotypic data only. In each case, clades usually retrieved as the sister group to amphisbaenians (lacertids for molecular data and dibamids for phenotypic data) were found within amphisbaenians (Supplementary Figs. 1-3). Contrary to the results recovered using a previous version of this data set^17^, we find considerable agreement concerning early diapsid relationships between total evidence non-clock and clock trees with results from Mr. Bayes (Supplementary Figs. S3-5).

In all of our results from total-evidence relaxed clocks, inferred rates of phenotypic evolution have their medians and means similar to each other, with modal, median and mean values ~2.0 for early evolving diapsid lineages during the Permian up to the end of the Middle Triassic (Fig. 1a, 2). In lepidosaurs, phenotypic and molecular rates have similar distributions, and median, mean and modal values between 0.3 and 1 (Fig. 1b, 3). Further, in lepidosaurs there is no detectable correlation between phenotypic and molecular rates (Fig. 1c), demonstrating a strong decoupling between both rates (supporting the utilization of separate clocks for phenotypic and molecular data herein). Among the periods of elevated rates of molecular evolution, only the early part of the Jurassic is coincident with relatively fast rates of phenotypic evolution. But even in this case, the branches exhibiting fast phenotypic change are not the same undergoing fast molecular change (Figs. 3 and 4).

**Fig. 1.**
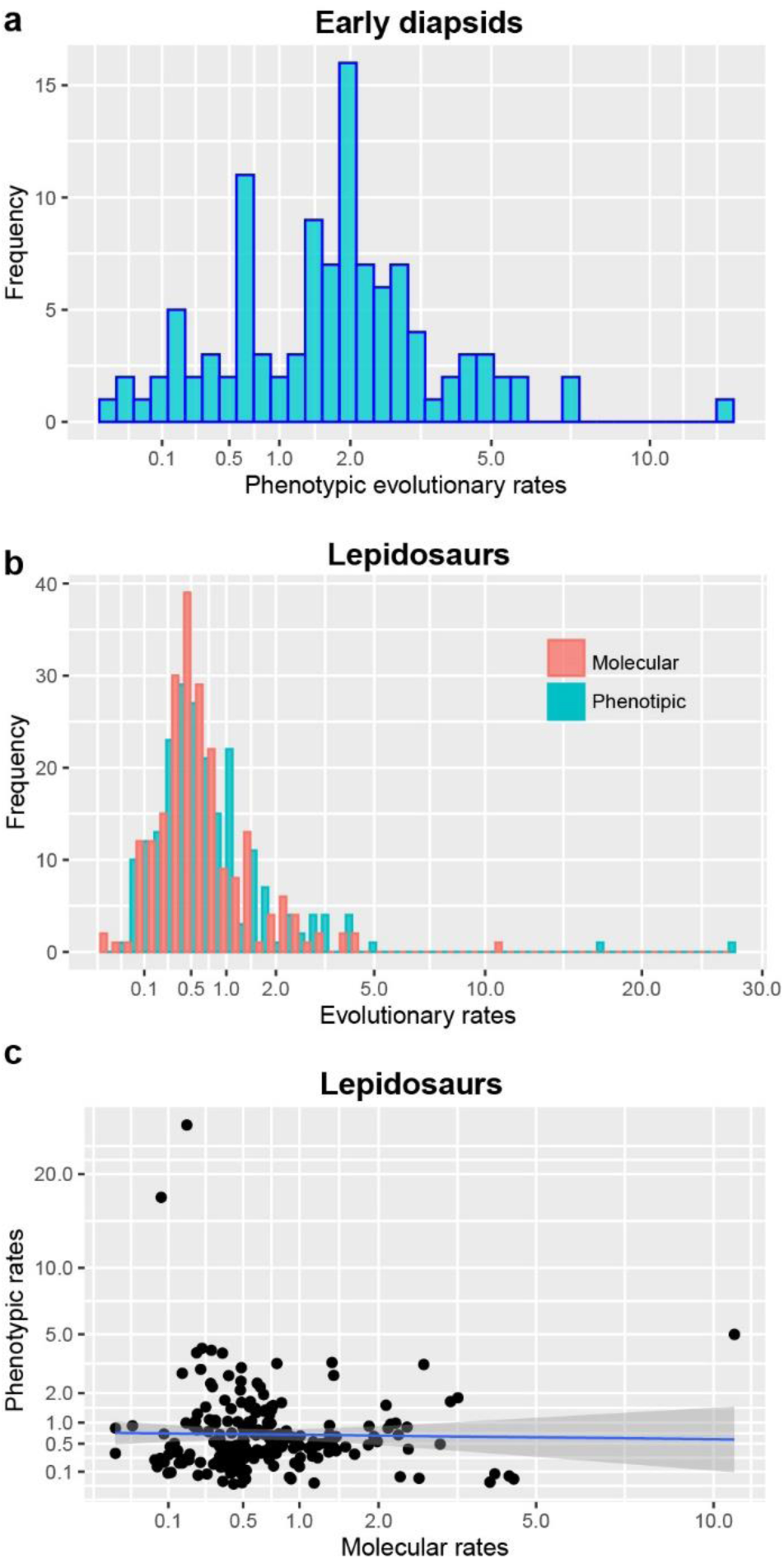
Distribution of phenotypic and molecular rates in different reptile groups. **A**, distribution of phenotypic rates among early evolving diapsid lineages (median=1.9; mean=2.1). **b**, distribution of phenotypic (median=0.51; mean=1.0) and molecular rates (median=0.54; mean=0.83) among lepidosaurs. **c**, linear regression between phenotypic and molecular rates (R-squared: 0.0001425; p-value: 0.8615) among lepidosaurs.

**Fig 2.**
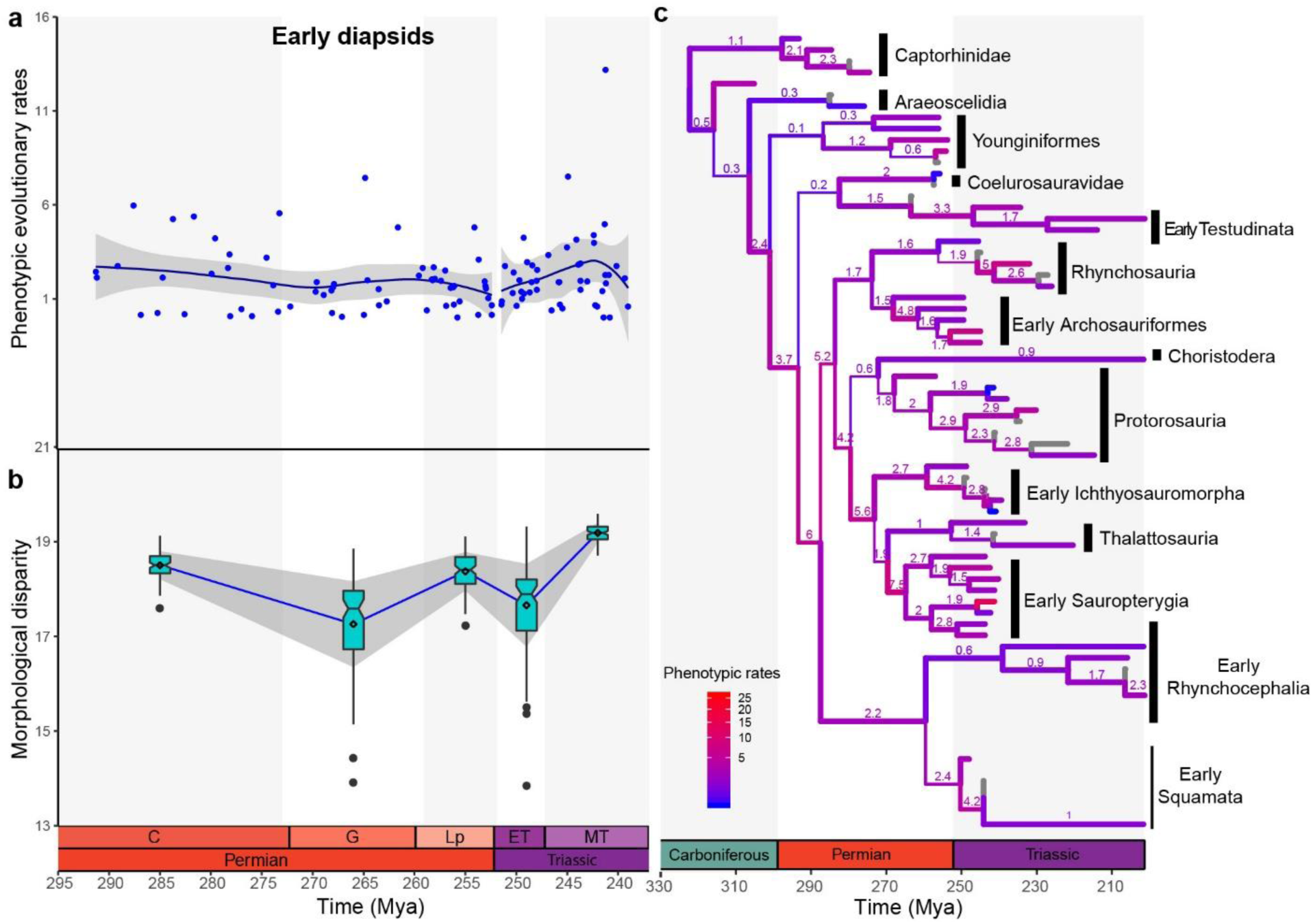
Phenotypic evolutionary rates and disparity through time in early diapsid reptiles. **a**, phenotypic rates among the major early evolving diapsid reptile lineages from the Early Permian to the Middle Triassic obtained from all unique bipartitions from the posterior trees. LOESS-smoothing trendline represent evolutionary rate fluctuations through time; grey area represents 95% confidence interval. Carboniferous rates are not considered here due to low sample size and extremely large confidence intervals. B, phenotypic disparity in early diapsids through time. Box plots represent distribution of 100 bootstraps at each time bin, with notching around the median and diamond indicating means. Blue trendline passes through median values and grey area represents one standard deviation. **c**, phenotypic rates of evolution in reptiles plotted on the time-calibrated maximum compatible tree from Mr. Bayes. Branch width proportional to posterior probabilities and branch values represent absolute phenotypic rates (character change per million year). For full tree see Supplementary Information.

**Fig 3.**
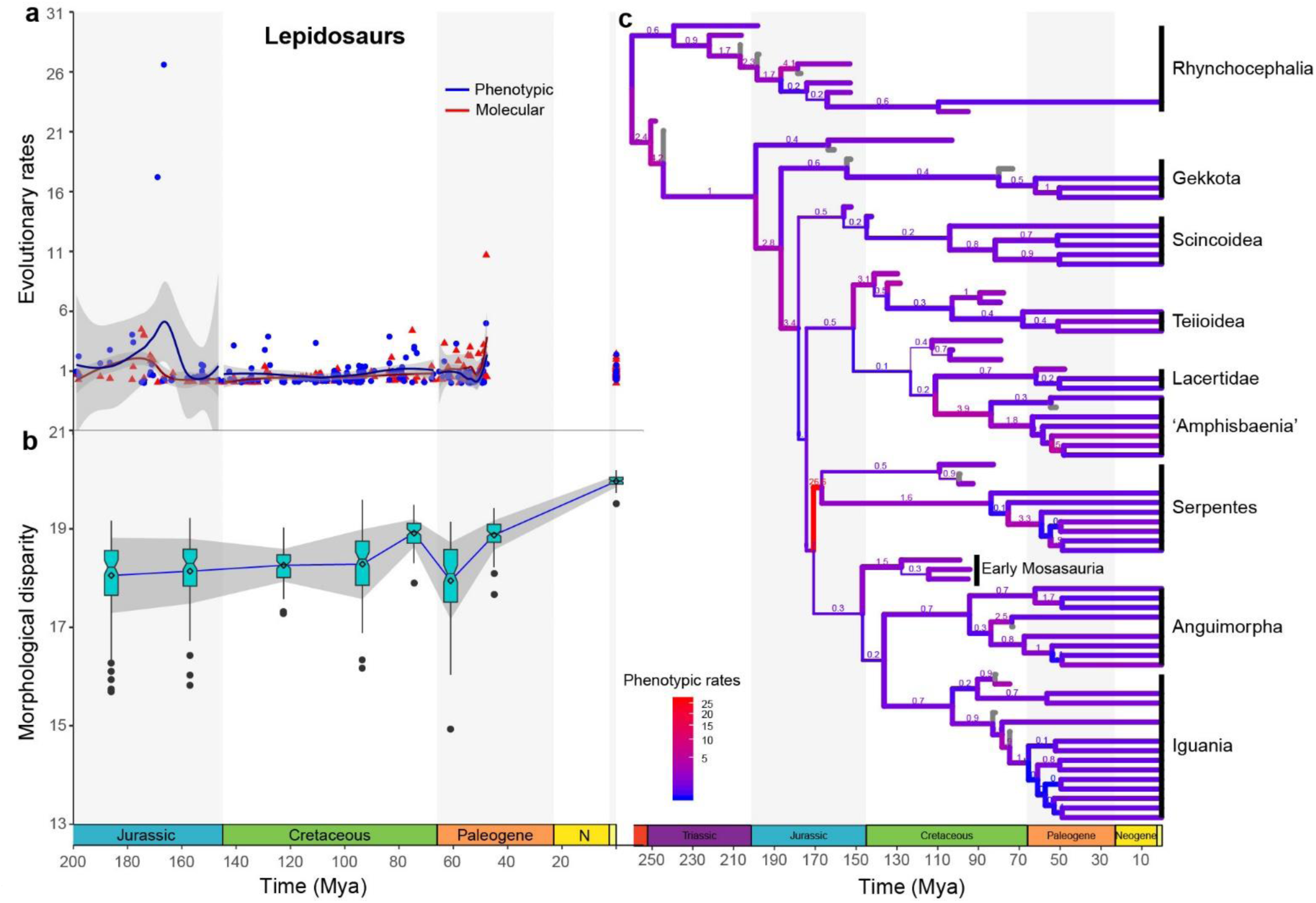
Phenotypic and molecular evolutionary rates and disparity through time in lepidosaurs. **a**, phenotypic and molecular rates among the major lepidosaur lineages from the Jurassic to the present time obtained from all unique bipartitions from the posterior trees. LOESS-smoothing trendline represent evolutionary rate fluctuations through time; grey area represents 95% confidence interval. Triassic rates are not considered here due to low sample size and extremely large confidence intervals. **b**, phenotypic disparity in early diapsids through time. Box plots represent distribution of 100 bootstraps at each time bin, with notching around the median and diamond indicating means. Blue trendline passes through median values and grey area represents one standard deviation. **c**, phenotypic rates of evolution in reptiles plotted on the time-calibrated maximum compatible tree from Mr. Bayes. Branch width proportional to posterior probabilities and branch values represent absolute phenotypic rates (character change per million year). For full tree see Supplementary Information.

**Fig 4.**
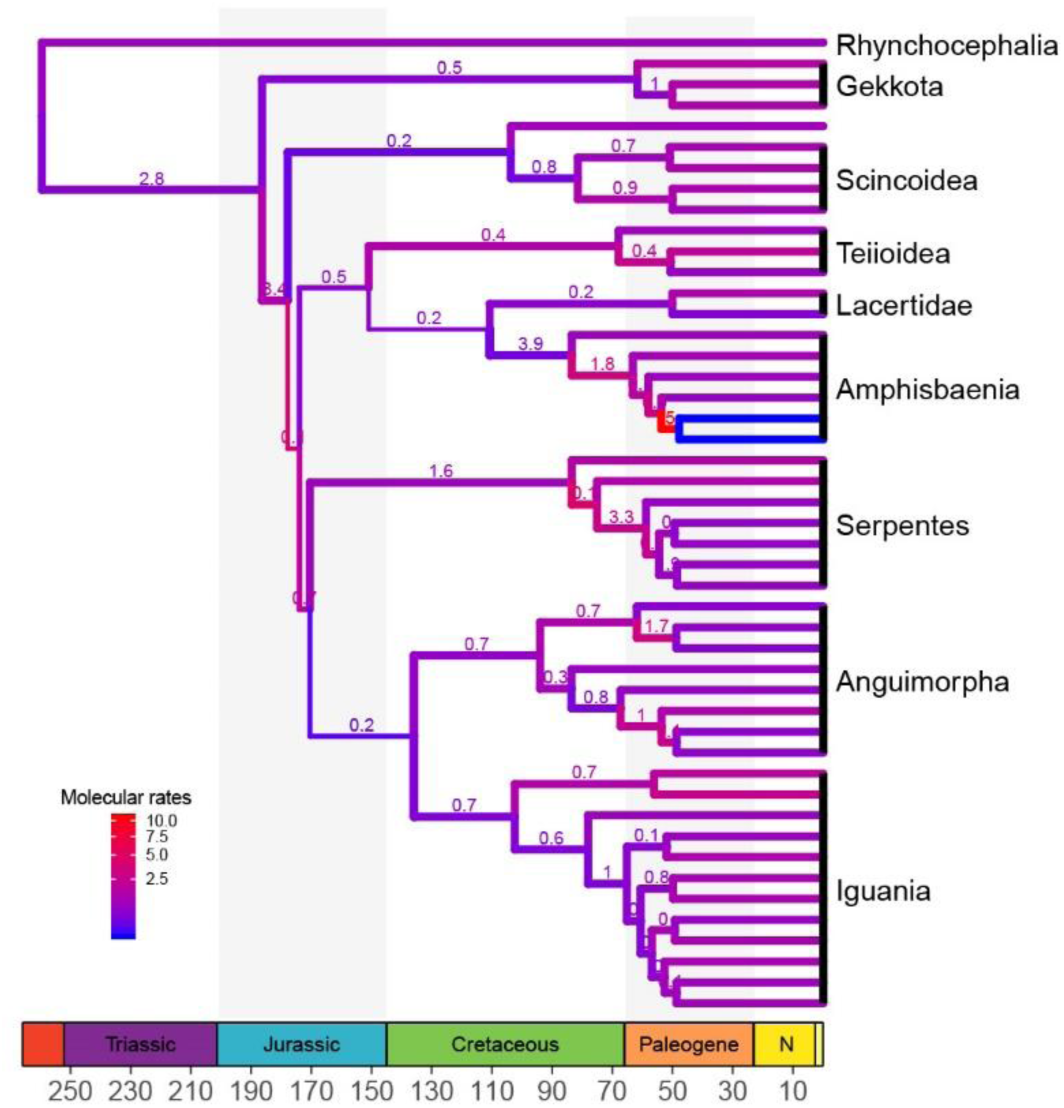
Molecular rates of evolution in lepidosaurs plotted on the time-calibrated maximum compatible tree. Molecular rates are plotted for all sampled extant taxa and internal branches until their most recent common ancestor. Branch width proportional to posterior probabilities and branch values represent absolute molecular rates (substitutions per million year). For full tree see Supplementary Information.

When observed across time, phenotypic rates of evolution in early diapsids are consistently accelerated (above one^26^ and well above modal values) during most of the Permian, indicating elevated rates of evolution at the origin of the major lineages of diapsid reptiles (Fig. 2a,c). This is coupled with relatively high rates of phenotypic disparity (Fig. 2b), although this disparity drops during the Guadalupian, and increases again during the Late Permian. It is difficult to be precise when most of the disparity was lost during the Guadalupian, but the early Guadalupian depicts some of the lowest phenotypic evolutionary rates during the Permian, which then increase at the Guadalupian-Lopingian transition. The disparity results suggest that some level of extinction followed by recovery of phenotypic diversity happened during the Permian, and thus during early diapsid history. Taxonomic diversity of terrestrial non-flying tetrapods has been recently demonstrated to also drop during the Guadalupian, followed by an increase by the end of the Permian^27^. These results support recent hypotheses that early diapsid reptiles were affected by the more recently discovered Guadalupian mass extinction^28^. However, the increase in evolutionary rates at the Guadalupian-Lopingian boundary is much milder compared to the increase in phenotypic disparity, thus deviating from the expected signal from a recovery event from a mass extinction. We note that the amount of data available for this particular timeslot concerning early reptiles in the present data set may not be enough to fully capture a shifting evolutionary rate regime at a high scale of resolution. Future addition of Middle Permian reptiles to our data set will provide a stronger assessment of shifts in evolutionary rates and its relationship to shifts in phenotypic disparity and taxonomic diversity, and assessing the impact of the Guadalupian mass extinction in early diapsids.

Phenotypic rates decrease at the Permian-Triassic boundary, but rapidly increase during the first few million years of the Triassic, reaching their peak at the Middle Triassic, subsequently starting to decrease at the end of the middle Triassic (Fig. 2a). Phenotypic disparity drops during the Early Triassic but recovers quickly and expands above pre-extinction levels by the Middle Triassic (Fig. 2b). These fluctuations in evolutionary rates and disparity levels match the expected patterns of an adaptive radiation in the aftermath of mass extinctions, in which the occupation of available ecological niches is associated with an expansion of phenotypic disparity and high evolutionary rates. This is further supported by the well-documented increase in the number of diapsid species and clades during the Triassic^24^.

In lepidosaurs (Figs 3), the beginning of the Jurassic marks the divergence of some of the deepest branches among major squamates clades, with elevated rates of phenotypic and molecular evolution at the origin of those clades as well as moderately high rates of phenotypic disparity. In general, those rates are lower compared to the rates observed at the origin and radiation of the major diapsid lineages during the Permian and Triassic. One major exception to the later trend, however, is the extremely high rate of phenotypic evolution at the origin of snakes. The branch leading to snakes is inferred to have the highest rates of phenotypic evolution among all the lineages of diapsid reptiles studied here (Figs. 2,3). High phenotypic rates during the early evolution of snakes were also recently found by another study based on extant snake taxa in a lepidosaur cranial shape data set ^29^. Elevated rates of phenotypic evolution suggest an additional potential explanation for the difficulty in estimating the phylogenetic placement of snakes among squamates using phenotypic data only, besides the issue of multiple independent cases of the reduction of limbs among lizards. Interestingly, molecular rates of evolution on the lineage leading to snakes are not as elevated, although they become higher within snakes, especially when compared to molecular rates of other squamates lineages (Fig. 3,4).

Both phenotypic and molecular evolutionary rates among lepidosaurs stabilize during the Cretaceous, lying closer to modal levels (Fig. 3a,b), whereas phenotypic disparity increased slowly but steadily throughout the Cretaceous, reaching its highest peak during the Mesozoic at the end of the Cretaceous (Fig. 3b). This gradual increase in phenotypic disparity is supported by the fossil record, as the Late Cretaceous sees the first appearance and subsequent increase in phenotypic disparity and taxonomic diversity, of the aquatically adapted mosasaurians^30^, appearance of the oldest preserved legged snakes^31^, along with the appearance of a large diversity of many crown group lizards in the Campanian and Maastrichtian of Mongolia^32^.

Lepidosaur phenotypic disparity drops following the Cretaceous-Paleogene mass extinction (Fig. 3b). Disparity, as well as phenotypic and molecular evolutionary rates (Fig. 3,4) remain relatively low during the Paleocene, with disparity increasing again during the Eocene, reaching pre-extinction levels. Phenotypic evolutionary rates do not see an equivalent increase to those observed for phenotypic disparity, although molecular rates are higher during the middle Eocene compared to earlier parts of the Paleogene. We lack sufficient data to estimate evolutionary rates during the Neogene, but rates among extant species remain relatively low while disparity is the highest in the history of lepidosaurs. The latter is supported by considering that squamates (essentially almost all of the extant diversity of lepidosaurs) comprise more than 10,000 living species, a level of taxonomic diversity that is inferred to be the highest in the history of the group^27^, including ecologically diverse forms inhabiting almost any environment outside of the polar circles.

## Discussions

Adaptive radiations are traditionally believed to be responsible for the origin of most of Earth’s taxonomic and phenotypic diversity, usually associated with the first stages of the evolution of major clades^1,4^. However, in our results, we only detected one instance in the history of early diapsid reptiles in which phenotypic evolution seems to have been driven by an adaptive radiation—the recovery from the aftermath of the PTME during the Triassic. At other periods of time we see patterns that are better explained by other models of evolution. For instance, we observe an important macroevolutionary lag between the time of origin (and initial phenotypic radiation) of the major diapsid lineages during the Permian and their later taxonomic diversification into several species within those major clades during the Triassic. This deviates from the classical model of adaptive radiation and instead matches the expectations of a pattern of early “disparification” without taxonomic richness (*sensu* ^33^) that is quickly followed by periods of loss and recovery of phenotypic disparity during the Late Permian.

Among lepidosaurs, we detected some of the highest rates of evolution during the early part of the Jurassic, at the time of diversification of some of the deepest branches in squamate evolution and the origin of important new body plans (e.g. snakes), but marked by little taxonomic diversity^27^. In contrast, both phenotypic and molecular evolutionary rates were stable and at low levels during the Cretaceous, the period where most of the first examples of modern lineages of squamates show up in the fossil record: the peak of Mesozoic taxonomic richness is reached at the end of the Cretaceous^27,34^. Additionally, overall phenotypic disparity slowly increased between the Jurassic and the end of the Cretaceous, indicating a slow and steady buildup of phenotypic space. Such a substantial gap of 100 million years between initially high rates of evolution and the much later acquisition of taxonomic richness, associated with a continuous construction of morphospace, is better characterized by the more recently proposed constructive radiation model^15^ that predicts that emergence of phenotypic novelties predate their taxonomic diversification by several millions of years. A major similar example is the fast evolution of phenotypic novelties and the exploration of morphospace during the early evolution of metazoans at the Cambrian explosion, generating many clades with few species, but with actual taxonomic diversification occurring much later in the history of animals^14,15^. This model notably, and importantly, departs from the classical adaptive radiation model that Simpson believed to be the predominant one governing megaevolutionary dynamics^1^.

In all of our results, phylogenetic branches with the highest phenotypic rates are frequently those within the early branches of newly evolving clades with markedly distinct new adaptive anatomical features characterizing new body plans (e.g., the emergence of turtles, marine reptiles, archosaurs, and snakes [Figs 2,3]). Early fast evolving branches in the history of new major clades were long predicted by Simpson (termed “tachytelic” lineages^1^), but were supposed to occur at periods of adaptive radiation. However, many bursts in phenotypic evolution are not observed here at times that can be characterized as adaptive radiations. Instead, we detected multiple bursts of phenotypic evolution throughout reptile history, as similarly observed during echinoid evolution ^11^. In contrast, high rates are also observed during the acquisition of unique body plans that represent “failed” evolutionary experiments (lineages that did not reach high levels of diversification and went extinct soon after their origin) such as placodonts during early sauropterygian evolution (Figs 2).

Surprisingly, clades with very similar functional adaptations exhibit radically different rates of phenotypic evolution. For instance, protective/armored morphotypes (turtles and placodonts), aquatic morphotypes (ichthyosaurs, thalattosaurs, eosauropterygians and mosasaurians), and serpentiform morphotypes (snakes and amphisbaenians) show highly distinct rates of evolution in their early history (Figs. 2,3), during the acquisition of their respective key phenotypic innovations. While the fastest evolving branch in early turtle evolution has rates up to 2.15 times faster than median values for overall rates of phenotypic evolution, the fastest branch in placodonts is 8.3 times faster than the median for early diapsids. The most dramatic example is represented by the extremely similar but convergent morphology of amphisbaenians and snakes (Fig. 5), which have markedly different rates of evolution at their origin (5 times faster than median values for lepidosaurs in amphisbaenians vs 34.1 times faster in snakes). To our knowledge, this is the first time that such levels of evolutionary rate heterogeneity among convergently evolving body plans has been detected. We note that, although usually characterized by the limblessness of its extant representatives, early snakes still retained partially developed limbs31, and many of the character changes contributing to those fast evolutionary rates relate to changes in the skull of both snakes and amphisbaenians, not limb evolution. Indeed, fast rates of skull shape evolution on the branch leading to snakes were found recently by^29^.

**Fig 5.**
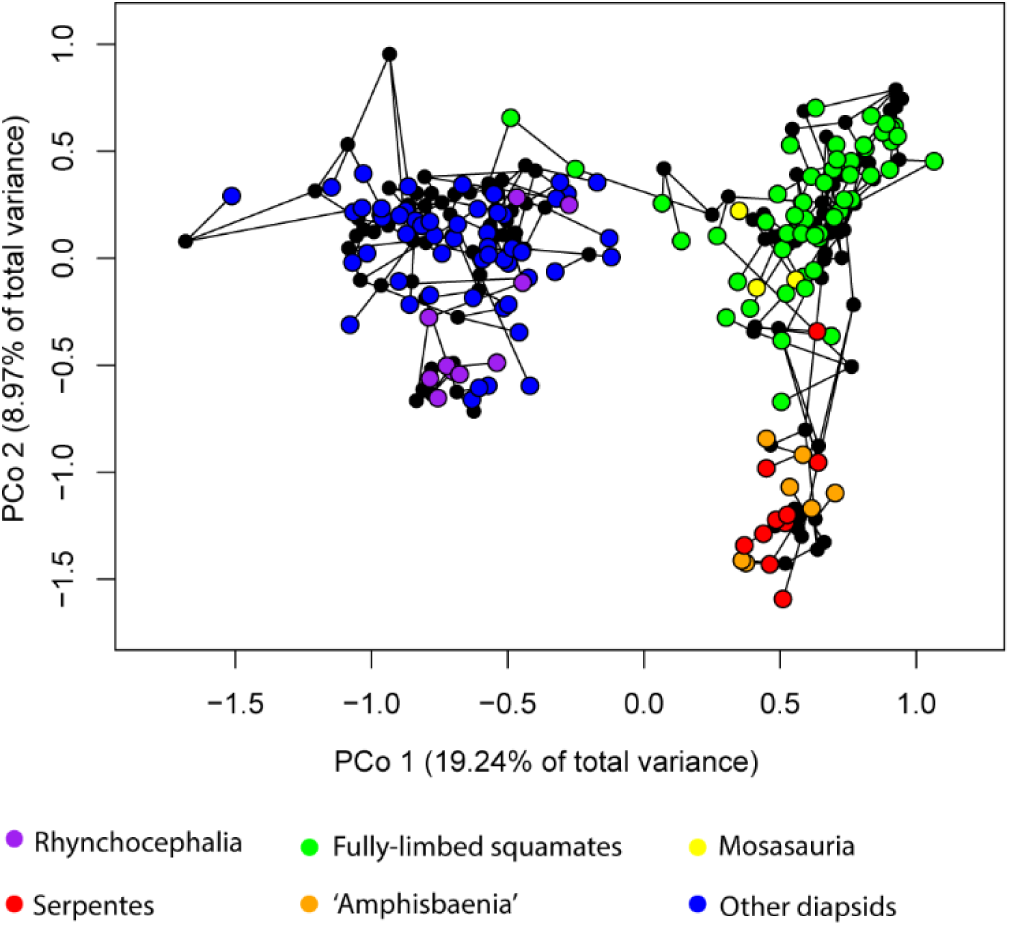
Phylomorphospace of diapsid reptiles. The first two axes of phenotypic variation among principal coordinates. Early diapsid reptiles occupy a distinct region of the morphospace from lepidosaurs, which in turn, have rhynchocephalians, non-serpentiform squamates, and serpentiform squamates (snakes and amphisbaenians) occupying different regions of the morphospace defined by PCo 1 and 2.

Contrary to the fast changes observed on phenotypic evolutionary rates, most deep nodes in lepidosaur evolution are marked by comparatively slower rates of molecular evolution (Fig. 1c,3,4). Interestingly, rates of molecular evolution are quite low on the branch leading to snakes and other clades marked by high levels of phenotypic evolution. Among the few empirical studies comparing phenotypic and molecular rates of evolution across broad time scales, a similar pattern is observed in mammals (despite using different methodologies), with molecular rates kept at relatively low levels during the early evolution of placental mammals^6^. Those results indicate that the structural protein coding sequences tested herein for lepidosaurs, and also in the mammalian study, do not seem to have any detectable correlation to the substantial and fast phenotypic changes observed at the origin of new body plans in diapsid reptile and mammalian evolution.

Such decoupling of evolutionary rates between phenotypic and protein coding sequences at the origin of major clades and new body plans provide empirical support for recent hypotheses concerning the genetic basis for major phenotypic changes at large evolutionary timescales. Although early theories on the genetic drivers of major phenotypic changes invoked revolutions in protein coding gene frequencies^1,35^, genomic studies have revealed that substantial phenotypic change appears to be mediated by changes on cis-(upstream) regulatory elements (CREs), and not by developmental gene duplication, or functional protein changes by mutations in coding sequences^36–38^. Therefore, our results from rates on protein coding sequences compared to rates of phenotypic evolution suggest that most genomic change associated with major phenotypic transitions in lepidosaurs (and possibly extinct reptile lineages that cannot be sampled for molecular data) might be located on conserved regulatory regions, as recently detected in the evolution of paleognathous birds^37^. Although outside the scope of the present study, we consider that the assessment of rates of evolution on conserved regulatory regions to be a fundamental next step on the investigation of the genomic basis for fast phenotypic change in reptiles.

Our results indicating exceptionally high phenotypic evolutionary rates at the origin of snakes further suggest that snakes not only possess a distinctive morphology within reptiles^39^, but also that the first steps towards the acquisition of the snake body plan was extremely fast. Therefore, snakes may hold some important goals towards understanding the processes driving phenotypic innovation in lepidosaurs. Some of such potential drivers may be represented by transposable elements (TEs). TEs can be found in large numbers on protein coding, intronic and regulatory sequences, eventually becoming exapted to novel functions, including regulation of gene expression within CREs in mammals^40^. The situation is even more dramatic in squamates, in which *Hox* gene clusters (which usually lack TEs and are conserved in most vertebrates to preserve the regulation of organismal development^41^) have an unparalleled accumulation of TEs compared to other vertebrates^40,42,43^. Transposable elements also have a role in the expansion of the number of microsatellites by microsatellite seeding^44^. In conformity with their large number of TEs, squamates have undergone microsatellite seeding during their evolution, and as a result, squamates have the highest abundance of microsatellites among vertebrates, with snakes, in particular, having the highest microsatellite content among eukaryotes^44^. It is therefore possible that the unusually high number of TEs and microsatellites in squamates (snakes in particular), and their subsequent exaptation to novel functions in the genome (both in protein coding and regulatory regions) may be one of the fundamental drivers of phenotypic innovation in squamates^40,42^, explaining the exceptional rates of evolution observed in snakes.

The patterns and processes governing the origin of major clades and new morphotypes across the tree of life remain poorly understood. Our study is one of the very few assessing such megaevolutionary dynamics over broad taxonomic and chronological scales, revealing a limited role of adaptive radiations at the origin of the major diapsid reptile clades and body plans. Although reptile evolution shows the classic signatures of an adaptive radiation following the PTME, we also detected fast evolving lineages and expansion of phenotypic disparity at periods of time not marked by adaptive radiations, as well as rate heterogeneity during the early evolution of similar morphotypes. How generalizable these patterns are across other major metazoan lineages remains to be determined. However, our findings lend support to the more recently proposed alternative models for the radiation of major lineages^15^, and hint at a more complex scenario concerning the evolution of reptiles in deep time than previously thought.

## Methods

Most paleobiological studies assessing rates of phenotypic evolution have utilized parsimony inferred phylogenetic trees for an *a posteriori* estimate of changes along the branches of the tree [e.g.^5,45,46^]. Those estimates have provided valuable insights into evolutionary dynamics in deep time, especially concerning fossil lineages, and to the understanding of detailed patterns of phenotypic change. However, an essential limitation of this approach is how to timescale the tree and the fact that frequently used parsimony trees minimize the number of changes along the branches, with both factors directly affecting rate estimates^47^. The integration of both phenotypic and molecular clocks in total evidence dating provides a powerful approach in which tree topology, divergence times, and phenotypic and molecular evolutionary rates, are jointly estimated, thus circumventing those limitations^47,48^. Further, estimates of divergence times and evolutionary rates can be averaged across the posterior sample of trees so that estimates take phylogenetic uncertainty into consideration. When those parameter estimates are taken directly from the Bayesian summary/consensus trees, those trees can be constructed based on the product of posterior probabilities or by selecting the posterior tree with highest posterior probability thus producing fully resolved trees upon which macroevolutionary parameters can be inferred without ambiguity, as is often the case with consensus trees derived from maximum parsimony (e.g., choosing between *acctran* vs. *deltran* approaches at polytomic nodes, or arbitrarily resolving polytomies). Additionally, different types of relaxed clock models are available (representing essentially distinct modes of evolution)^49,50^, and can be tested in order to determine which one has the better fit to the data set, thus allowing an essential simultaneous consideration of tempo and mode towards estimating evolutionary relationships and rates of evolution.

### Morphological and molecular data sets

Here we updated the recently published diapsid-squamate data set of Simões *et al.*^17^ in order to expand the representativeness of extant taxa, which are informative on both morphological and molecular data. Twelve additional taxa were added to this data set: nine extant species (the snakes *Rena humilis, Afrotyphlops punctatus, Python regius* and *Lichanura trivirgata*, the amphisbaenians *Amphisbaena alba, Trogonophis wiegmanni*, and three additional limbed lizards, *Tupinambis teguixin, Celestus stenurus* and *Varanus albigularis*) and three fossil taxa (*Pleurosaurus_goldfussi, Gobiderma pulchrum* and *Cryptolacerta hassiaca*). Morphological data was collected for the additional taxa based on personal observations (by T.R.S.) and molecular data from the nine extant taxa were added to the molecular component of this data set^17^. Three taxa that operate as wildcards in the present data set, as identified in a previous study using the RogueNaRok algorithm^17,51^, were removed for the present analyses, namely *Paliguana whitei*, *Palaeagama vielhaueri*, and *Pamelina polonica*, resulting in considerable improvement on convergency of resolution of early diapsid relationships between non-clock and clock trees (see Results).

The molecular data set for the selected coding regions were obtained from GenBank (Supplementary Table S1, Supplementary Data 2). For *Python regius*, for which molecular data were not available, we used sequences of congeneric species, *P. molurus*. Sequences were aligned in MAFFT 7.245^52^ online server using the global alignment strategy with iterative refinement and consistency scores (G-INS-i). Molecular sequences from all extant taxa were analyzed for the best partitioning scheme and model of evolution using PartitionFinder2^53^ under Bayesian information criterion (BIC).

### Bayesian inference analyses

Both non-clock and clock based Bayesian inference analyses were conducted using Mr. Bayes v. 3.2.6^18^ and the BEAST2 package^19^ using high performance computing resources made available through Compute Canada. Molecular partitions were analyzed using the models of evolution obtained from PartitionFinder2^53^ (see dataset), and the morphological partition was analyzed with the Mkv model.

### Time-calibrated relaxed clock Bayesian inference analyses

We implemented “total-evidence-dating” (TED) using the fossilized birth-death tree model with sampled ancestors (FBD-SA), under relaxed clock models in Mr. Bayes v.3.2.6^21,54^—100 million generations, with four independent runs with six chains each, and a gamma prior of rate variation across characters. We conducted the same analysis using the BEAST2 package^19^, with four independent runs, also with a gamma prior of rate variation across characters. To ensure that each independent single chain run in BEAST2 reached stationarity, we increased length of each analysis to 200 million generations. Runs were sampled every 500 generation with the initial 55% of samples removed as ‘burn-in’. We provided an informative prior to the base of the clock rate based on the previous non-clock analysis: the median value for tree height in substitutions from posterior trees divided by the age of the tree based on the median of the distribution for the root prior: 23.8582/325.45 = 0.0733, in natural log scale = −2.61308. We chose to use the exponent of the mean to provide a broad standard deviation: *e*^0.0733^ =1.076053. The vast majority of our calibrations were based on tip-dating, which accounts for the uncertainty in the placement of fossil taxa and avoids the issue of constraining priors on taxon relationships when implementing bound estimates for node-based age calibrations54. The range of the stratigraphic occurrence of the fossils used for tip-dating here were used to inform the uniform prior distributions on the age of those same fossil tips (thus allowing for uncertainty on the age of the fossils).

Convergence of independent runs was assessed using: average standard deviation of split frequencies (ASDSF ~ 0.01), potential scale reduction factors [PSRF ≈ 1 for all parameters] and effective sample size (ESS) for each parameter was greater than 200 for Mr. Bayes. Independent BEAST runs were combined using LogCombiner v2.5.1 (available with the BEAST2 package) and checked for stationarity and convergence in Tracer v. 1.7.155. The ESS value of each parameter was greater than 200 and ASDSF < 0.01.

### Testing for the best fitting clock prior

Distinct clock models allow for different assumptions regarding the predominant tempo and mode of evolution. While strict clocks presume constant rates of evolution across lineages, relaxed clock models allow for changes in the rate of evolution among lineage. For instance, relaxed clocks include models where rates at each branch in a phylogeny is drawn independently and identically from an underlying rate distribution (uncorrelated clocks), to others where the rate at a particular branch is dependent on the rates on the neighbouring branches (autocorrelated clocks)—see^49,50^ for modelled comparisons. Therefore, in uncorrelated clocks rates are free to change more dramatically among neighbouring branches, resulting in shorter branch lengths and higher rates than autocorrelated clock models^54^, and thus reflecting a more punctuated model of evolution compared to the more gradualistic model represented by autocorrelated rates^50^.

In order to detect the most appropriate clock models, we used Bayes factors (BF) applying model fitting analyses using the stepping-stone sampling strategy to assess the marginal model likelihoods^56^ for each clock model for the current data set (50 steps for 100 million generations in Mr. Bayes and 100 million generations in BEAST2 (two runs each). We tested between strict clock models and relaxed clock models. Relaxed clock models were further tested for linked clock models (where morphological and molecular partitions share the same clock, and therefore variations on the rate of evolution) and unlinked clock models, allowing for the clock rates to vary independently among the morphological and molecular partitions of the data set. In Mr. Bayes, we found a considerably stronger fit for relaxed clock models against a strict clock model (BF>2000), and a stronger fit (BF >400) for unlinked clock models, thus supporting the treatment of morphological and molecular rates independently. The results are expected given the broad scale of the present data set, which is inclusive of several reptile families sampled over the last 300 million years and characters from multiple regions of the phenotype and genotype. Finally, an independent gamma rate (IGR) unlinked relaxed-clock model was favoured relative to the autocorrelated clock model (BF =150), indicating the data supports a model allowing more disparate shifts in evolutionary rates across lineages.

The stepping-stone analyses conducted in BEAST2 ran for 100 million generations, with some of the longest runs (using random local clocks) taking 37 days to complete in a computer cluster, and yet they failed to reach the stationarity phase. The large taxonomic sampling of our data set (which increases computational time exponentially) compared to most other total evidence data sets analysed under BEAST2 indicate that the sheer size of this data set prevents a reasonable assessment of marginal likelihoods independently from the ones performed under Mr. Bayes. Therefore, for subsequent analyses using BEAST2 we implemented the two uncorrelated relaxed clock models available in BEAST2 (lognormal and exponential), given the much stronger fit of uncorrelated relaxed clock models over other clock models in Mr. Bayes. Further, relaxed clock implementations can recover homogeneous rates of evolution when the best fit model is supposed to be clock like^57^, indicating relaxed clock models can fit a variety of different evolutionary scenarios.

### Divergence time estimates and evolutionary rates

A “diversity” sampling tree prior is implemented in Mr. Bayes v. 3.2.6, but it is not yet available on BEAST2. Accounting for “diversity” sampling impacts tree priors ^20^, affecting divergence time precision and accuracy 21,54. The results from our initial relaxed clock analyses show considerably (and unreasonably) older divergence times from the trees using BEAST2 compared to Mr. Bayes (tens of millions of years older). Considering the main prior choices were the same between the two software packages, we attribute the much older divergence times in BEAST2 to not accounting for diversity sampling in total evidence analyses, as already demonstrated by previous studies ^21,54^. Additionally, factors such as vague priors, limitations on currently available models of morphological evolution as well as conflict between the morphological and molecular signal may result in pushing divergence times further back in time (exceptionally long ghost lineages), especially among the deepest nodes on broad scale phylogenies, contributing to the phenomenon of “deep root attraction” (DRA) ^22^. It is possible to minimize this impact by providing informative priors that decrease the likelihood of long ghost lineages, such as modelling higher diversification rates or low extinction probability. This correction for DRA could potentially provide a tool for correcting the overestimation of divergence times in BEAST2, as reported above. Following Ronquist *et al.*^22^, we implemented one of those strategies (specifically, giving higher probabilities of low extinction by placing a Beta (1,100) on the turnover probability prior) to assess its impact on divergence times on the analyses conducted on both Mr. Bayes and BEAST2.

Implementing this strategy highly increased the precision of divergence times among the oldest nodes on the summary tree from Mr. Bayes, and also brought divergence times for the oldest nodes on the tree into much greater agreement with the fossil record (see Results). For instance, in the analysis with no DRA correction average variance of divergence times among the 50 oldest nodes taken from the posterior trees was 143.07 million years (myr), whereas it was 58.88myr among the 50 youngest nodes (excluding extant nodes); in the analysis with DRA correction those respective values decreased to 25.3myr and 16.49myr (see also ranges of 95%HPD between those analyses in Supplementary Figs 4,5). Therefore, we used the results from the DRA corrected analyses to report divergence times and evolutionary rates. Notably, even accounting for low extinction probability to reduce DRA in our analyses using BEAST2, we noticed no visible difference in divergence times among the oldest nodes. Divergence times were still considerably older (frequently 10-20 million years older) among intermediary and older nodes in the maximum clade credibility tree compared to the summary tree from Mr. Bayes (Supplementary Figs 6,7). This suggests that informative tree priors are not enough to avoid overestimating divergence times when diversity sampling is not taken into account, at least in BEAST2.

Branch length estimates and tree calibration invariably impact estimates of absolute rate values and correlating rates with specific periods of time in the geological record is affected when divergence times are biased. As a result, our main results report only the trees from Mr. Bayes, where divergence times are not being overestimated by DRA.

### Data set adaptation for morphological disparity analyses

Phylogenetic morphological characters provide a large number of variables that can be easily utilized for morphospace analysis and have been implemented in a large variety of studies on different taxonomic groups. Importantly, discrete phylogenetic characters can easily capture the disparate morphological variation that is observed among higher taxa (as observed in broad scale phylogenies, such as in the present data set) [e.g. ^58^]. Yet, important adaptations and considerations of phylogenetic data sets need to be taken into account for such kind of analyses, as further described below.

Large amounts of missing data, usually above 25%, considerably reduce the overall distance between taxa that can be captured on ordination spaces on both empirical and simulated data sets ^47,59^. To reduce the negative impact of missing data, we removed all characters with more than 30% of missing data from the data set, which resulted in a total of 19% missing data on the final data set—safely below the threshold of 25% ^47,59,60^. Additionally, inapplicable characters are a big conceptual problem to construct a morphospace. Taxa with inapplicable characters will have their placement enforced upon a space they do not reside in (which is conceptually very different from missing data—when they reside in that space, but we currently lack data to place them) ^60^. Deleting all inapplicable characters would further decrease the number of utilized characters at about 30%, thus reducing the span of morphological representation in the data set. Therefore, to avoid inapplicable characters, but keep minimal representation, we deleted all characters that were inapplicable to more than 5% of taxa, rescoring the remaining cells as missing data. Polymorphisms were converted into NA scores (treated as “?” during analyses), following previous recommendations and based on the reasoning above ^58,60^.

Autapomorphies, if unevenly sampled across taxa, may also contribute to bias distance matrices. However, if autapomorphies are uniformly distributed across terminal taxa, then their overall effect is to increase overall pairwise distance between terminal taxa uniformly, therefore not creating biasing the interpretation of the data ^61^. In the present data set, there are some directly observed autapomorphies, which were already excluded during the removal of characters with large amounts of missing data (see below). Having no remaining autapomorphies in the data set is another way of having a uniform distribution of autapomorphies, guaranteeing that no taxon will have additional dimensions separating it from other taxa in the dissimilarity matrix and morphospace ordination procedure.

### Intertaxon distance matrix and ordination matrix

The procedures above resulted in a final reduced matrix of 138 taxa and 105 characters. This number of characters is more than sufficient to provide reasonable estimates of disparity ^62^. This data set (data set 1) was used to construct a morphospace for all sampled clades of reptiles. A second version of the data set (data set 2) was adapted to compare disparity across time. Since most post-Triassic taxa in the data matrix are lepidosaurs, we deleted the only three non-lepidosaurian post-Triassic taxa (*Kayentachelys*, *Philydrosaurus* and *Champsosaurus*), in order to distinguish disparity across time among early diapsids clades in general (between the Late Carboniferous and Late Triassic) and disparity across time among lepidosaurs (Early Jurassic to the present). Therefore, data set 2 contained 135 taxa and 105 characters (and the original time calibrated tree was also pruned of those three taxa to match the reduced data set).

Using the reduced data sets and the time calibrated trees, we constructed an intertaxon distance matrix **D** and an ordination matrix using principal coordinate analysis (PCoA— or classical multidimensional scaling). We implemented MORD as our method of estimating pairwise taxon distances for the distance matrix **D** and the subsequent ordination matrix, made available through the Claddis R package ^47^, implementing Cailliez’s correction for negative eigenvalues. Additionally, we increased our sample size by including internal nodes using ancestral state reconstructions through the recently developed pre-OASE1 method ^63^. This procedure provides a much better approximation of the true morphospace when compared to methods to reconstruct ancestral nodes in most previous disparity studies using ancestral state reconstructions ^63^.

### Morphological disparity measures

Here, we used the sum of the variances (a post-ordination metric), which is comparatively robust to sample size and is not affected by the orientation of the coordinate axes of the ordination analysis ^58,62^. Importantly, only post-ordination methods can be used to produce a morphospace projection. Further, post-ordination methods have been more widely used in the literature making our results more directly comparable to previous studies. Since PCo scores based on phylogenetic data usually have the first principle axis representing a small proportion of the total variance (usually the first two PCo representing less than 50% of total variance), morphospace representation using PCo scores should be taken with caution. This problem can be avoided in our assessment of disparity across time (our main measure of both chronological and taxonomic changes in disparity), in which it is possible to take into account all axes of variation to estimate morphological disparity. Nonmetric multidimensional scaling was not used to preserve the metric properties of the dissimilarity matrix. To measure morphological disparity across successive time bins, we used the R package dispRity 64 to subdivide the data across time bins. Our data is not evenly sampled across time, as it was designed to maximize taxonomic representation across stratigraphic intervals. Therefore, uniform time bins would create more heterogenic sample sizes across bins with some bins containing drastically low sample values, besides not capturing important geological boundaries reflective of important environmental shifts and mass extinctions. For those reasons, we chose time bins approximating stratigraphic intervals to subdivide our data chronologically, which enables capturing changes across major mass extinctions at stratigraphic boundaries (e.g. Permian-Triassic and Cretaceous-Palaeogene mass extinctions), and also less heterogenic sample sizes across the bins.

Additional methodological details can be found in Supplementary Methods in the Supplementary Information file. Supplementary Data S1-S5 (data sets in Nexus format) along with trees, log files, prior parameters and posterior parameter values described in the results and figures can be found online at: NNNNNNNNNNN

## Supplementary Information

is linked to the online version of the paper at NNNNNNNNN

## Acknowledgements

T.R.S. was supported by an Alexander Agassiz Postdoctoral Fellowship (Museum of Comparative Zoology, Harvard University). O.V. was supported by the Natural Science and Engineering Research Council of Canada (NSERC) Discovery Grant 327448 to Alison M. Murray and Alberta Ukrainian Centennial Scholarship. We also thank several curators that allowed us to have access to all the specimens analysed in this study.

## Author Contributions

T.R.S. led on manuscript writing, morphological dataset construction and conducted disparity analyses; T.R.S. and O.V. performed molecular sequence alignment and phylogenetic analyses; all authors contributed to discussions and manuscript editing.

